# Senescent widow spiders are poor mates but remain attractive to mate-seeking males through deceptive signalling

**DOI:** 10.64898/2026.03.04.709482

**Authors:** Andreas Fischer, Beatrice Chee, Andrea C. Roman Torres, Gerhard Gries

## Abstract

As fertility declines with age, unmated senescing female animals experience an increasing need to secure a mate. Elevated pheromone production and extended pheromone release are known mechanisms underlying deceptive sexual signalling in relation to the progressive age of signalers but both mechanisms incur elevated metabolic expenses. Here, we report an intricate new mechanism of deceptive signalling that allows unmated senescing female false widow spiders, *Steatoda grossa*, to conserve metabolic costs for silk and pheromone production while still achieving sustained attractiveness to mate-seeking males. Senescing females produced “honest” state-dependent signals, saving metabolic expenses by building webs with fewer silk strands and depositing less courtship-inducing contact pheromone on their web. However, senescing females concurrently engaged in deceptive signalling in that they remained as attractive to males as young(er) females by accelerating the hydrolytic conversion of web-borne contact pheromone components to air-borne mate-attractant pheromone components. Accelerated dissemination of mate-attractant pheromone from webs was correlated with an age-linked increase in web pH, which we posit enhanced the enzymatic activity of a web-borne carboxyl-ester-hydrolase. Essentially, senescing females concealed their low residual reproductive value by sustained high-level pheromone dissemination. Because the lifetime reproductive output of females declined with age at mating, old females are indeed poor prospective mates and thus deceptive signalers. Their deceptive mate-attractant signals seem evolutionary stable because they rarely remain unmated. Moreover, the males’ reproductive fitness benefit of mating with an old(er) female – and thus siring fewer offspring than mating with a young female – may still outweigh the costs of rejecting an old female and resuming mate search for a young female, a search which may not be successful.

## Introduction

Attraction of mates is essential for sexual reproduction. Consequently, diverse sexual signals have evolved across animal taxa^1^ that are based on various sensory modalities, including smell, taste, vision, audition, and somatosensation^2^. Of all signal modalities, chemical signals are deemed the most ancestral and widely-used across animals^3^.

Mate-attractant pheromones are airborne intraspecific chemical signals^4^ that attract potential mates from afar. Contact pheromones, in contrast, are sensed upon physical contact with the signaller^5^, revealing their identity and various physical characteristics. The information encoded in sex pheromones can be diverse, ranging from mate availability^6^ to resource access^7^, and may even reveal the signaller’s age^8^. Pheromones are generally considered cheap to produce^5,9–11^ but evidence to support this concept was reported in only a single study^12^. Challenging this concept, fitness costs such as shorter lifespans stemming from pheromone production have been demonstrated in four empirical studies^13–16^. Pheromone production may require to reallocate energy from essential physiological processes^17^. Ecological fitness costs associated with pheromone production and emission arise from predators eavesdropping on pheromones to locate prey^11,18^. Thus, sexual communication via sex pheromones presents a trade off between mate-attraction and survival^19^.

Strategic signaling evolved as a trade-off between signal costs and the need for mates^19^. Reproductive senescence – the age-linked decline in fertility and reproductive performance^20^ – presents animals with an ever-increasing need to secure sperm and to allocate resources to immediate reproduction rather than self maintenance (terminal investment concept)^21,22^. Consequently, old physically poor animals may produce ‘need-dependent’ dishonest signals that are highly attractive to signal recipients (Fig. 1a, orange segment). Conversely, old animals disseminate honest signals that truly reflect their poor physical state (Fig. 1a, pink segment).

**Figure 1.**
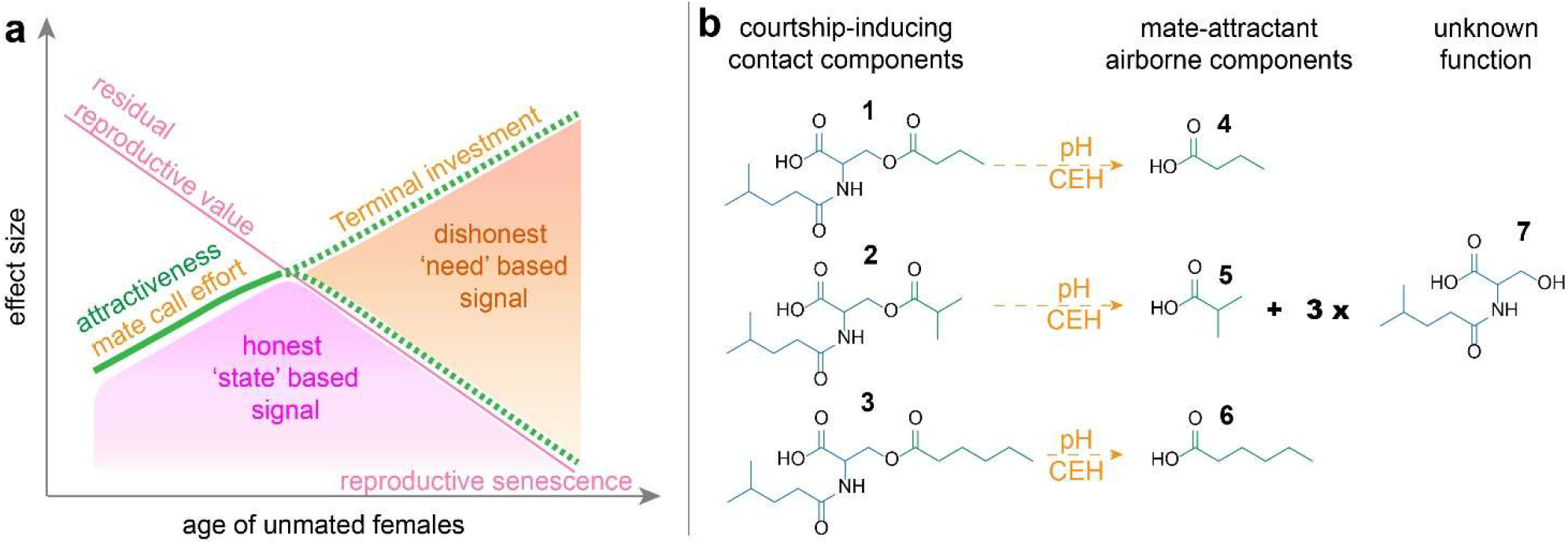
General signal theory applied to reproductive senescence and sexual pheromone signals of female false widow spiders, *Steatoda grossa*. (a) Theoretical concept of age effects of unmated females on their mate call investment (deposition of contact pheromone components and release rate regulation of mate-attractant pheromone components = web attractiveness), and the effect of female age at mating and female reproductive output. The dashed attractiveness line represents either a declining honest (‘state-based’) signal (pink area), or a deceptive (‘need-based’) signal (orange area). (b) Pheromone components of female *S. grossa*. The web-borne contact pheromone components [*N*-4-methylvaleroyl-*O*-butyroyl-L-serine (**1**), *N*-4-methylvaleroyl-*O*-isobutyroyl-L-serine (**2**), and *N*-4-methylvaleroyl-*O*-haxanoyl-L-serine (**3**)] induce courtship by males upon contact but hydrolyse at their ester bond, giving rise to airborne mate-attractant pheromone components [butyric- (**4**), isobutyric- (**5**) and hexanoic (**6**) acids], with the corresponding hydrolysis product *N*-4-methylvaleroyl-L-serine (**7**) accumulating on the web. The hydrolysis is thought be mediated by the pH-dependent activity of a carboxyl-ester-hydrolase (CEH).

Conflict between the sexes arises from deceptive signaling as a terminal investment strategy^22^. The signal recipient is deceived by being attracted to, and mating with, a signaler perceived to have high fitness but actually being reproductively senescent. Recipients of dishonest signals incur further costs related to mate-search^23–25^ and courtship^26,27^. Age-linked dishonest signalling seems widespread. A meta-analysis of 147 species across animal taxa found that old unmated males were consistent ‘need-based’ deceptive signallers^28^. Similarly, a review of 44 studies revealed that aging females of many moth species increased their efforts to attract a mate^8^.

Sexual signalling studies focused on males and offered little information on females^28–31^. In addition to this gender bias, there is a taxonomic bias favoring insects while neglecting other animal taxa such as spiders^32–34^.

Female web-building spiders disseminate airborne mate-attractant pheromone components from their webs that attract males^35^. Upon arrival on a web, the male senses silk-borne contact pheromone components which induce courtship by the male in a dose-dependent manner^36,37^. Male spiders sense pheromones via specialized chemo-sensilla on their legs, wall-pore sensilla for airborne pheromones and tip-pore sensilla for contact pheromones^38,39^. In the false widow spider, *Steatoda grossa*, the courtship-inducing contact pheromone components [*N*-4-methylvaleroyl-*O*-butyroyl-L-serine (**1**), *N*-4-methylvaleroyl-*O*-isobutyroyl-L-serine (**2**), and *N*-4-methylvaleroyl-*O*-haxanoyl-L-serine (**3**)] hydrolyse at their ester bonds and give rise to mate-attractant pheromone components [butyric-(**4**), isobutyric- (**5**) and hexanoic (**6**) acids] (Fig. 1b)^37^ that disseminate from the web, whereas the hydrolysis product *N*-4-methylvaleroyl-L-serine (**7**) accumulates on the web (Fig. 1b). This hydrolytic pheromone conversion is thought to be catalyzed by the pH-dependent activity of a carboxyl-ester-hydrolase^37^. Because females can manipulate their webs’ pH^40,41^, and thus the rate of pheromone hydrolysis, they can adjust the dissemination of mate-attractant pheromone and thereby the attractiveness of their webs^42,43^.

Female *S. grossa* are long-lived (>350 days; suppl. Fig. 1) global synanthropic spiders that commonly inhabit buildings ^44,45^ where males may be scarce. Considering the females’ long life and known sexual communication system, we chose *S. grossa* as a model organism for our study, testing for senescence-related deceptive signalling by females and investigating the underlying chemical mechanisms.

We tested three hypotheses (H): senescing females (1) produce progressively fewer offspring; (2) adjust their web density and sexual pheromone signals in accordance with their diminishing metabolic resources; and (3) attract males as effectively as young females through deceptive signalling.

## Methods

### Spider maintenance

Experimental *S. grossa* were the F2 and F3 progeny of adult females collected around the insectary of Simon Fraser University (Burnaby Campus, Canada, 49°16’38.9” N 122°54’58.1” W)^6^. Spiders were fed weekly and raised individually in petri dishes (100 × 20 mm) until they reached sexual maturity. Mature females were then transferred to 300 mL plastic cups (Western Family, Tigard Oregon, USA). Juvenile spiders and sexually mature males were fed *Drosophila* vinegar flies, whereas female spiders received black blow flies, *Phormia regina*. All spiders had access to water in cotton wicks. Water and food were provided once per week. The body weight of unmated sexually mature females and males was measured using a TR-204 Denver Instrument Company Scale (Denver Instruments, Arvada, Colorado, USA). The size of adult spiders was approximated as the average tibia-patella length of the first leg pair^46^. The body condition of adult females was then determined as the residuals of the weight and size regression, whereas for males the log-transformed weight was used^47,48^. Spider age refers to the time since their maturity molt.

### Web collection

Each unmated female was allowed 72 h to build her web on a triangular prism frame (8.5 × 8.5 × 8.5 cm) built from bamboo skewer sticks (Good Cook, San Leandro, CA, USA)^48^. Each frame was set at the center of a tray filled with water to ensure that the female could not escape or come into contact with other females.

### H1: Senescing females produce progressively fewer offspring

The reproductive fitness of females was measured as their lifetime reproductive output of viable spiderlings. Forty-one unrelated, unmated females were selected from a pool of 605 females. Their apparent body condition was seemingly not correlated with age (GLM, χ^2^ = 0.55, df = 1, p = 0.457). Each female – 5 to 996 days old – was allowed to mate with an unrelated unmated 16-day-old male. The custom-built plexiglass ‘mating’ arena (46 cm × 17 cm × 21 cm) consisted of three chambers (15 cm × 17 cm × 21 cm each), each housing a female on her web-bearing frame to which a male was added. All female-male pairs were kept 8 h in a chamber, and their behaviour was recorded with a Sony Exmor R Camera (Sony, Tokyo, Japan). Videos were analyzed using VLC Media Player (VideoLAN, Paris, France) to determine the duration of courtship behaviour as well as the occurrence and duration of copulation^48^.

The lifetime reproductive output of each female (that was sustained as described above and mated once at a specific age) was measured by (*i*) removing each of her sequentially-produced egg sacs one week prior to spiderling emergence, (*ii*) placing the egg sac into a petri-dish with moist cotton, (*iii*) counting all spiderlings emerging from the egg sac, and (*iv*) tallying up the spiderlings from all egg sacs the female produced until she eventually died.

### H2: Senescing females adjust their web density and sexual pheromone signals in accordance with their diminishing metabolic resources

We quantified age-dependent sexual signaling of 70 female spiders that were 19 to 815 days old but had similar, age-independent, body condition (GLM, χ^2^ = 1.13, df = 1, p = 0.287). Each female was allowed to sequentially build two webs on a large frame (18 × 18 × 18 × 25 cm). The first web was used to quantify the silk strand density of the web, and the amount of contact pheromone deposited on the web, whereas the second web was used to determine the web’s pH.

Silk strand density of webs was determined following an established protocol^42,43^. Silk strands were counted along nine axes using a thin metal rod marked at 1-cm intervals that was inserted into the web. We made three vertical insertions at 1 cm from the three corners of the frame (H), three horizontal insertions at the top of the frame from each corner to the center of the corresponding hypothenuses (S), as well as three insertions parallel to the latter but half-way down the frame (G). Overall, we recorded the presence or absence of a silk strand at 22 and 15 1-cm intervals for each vertical insertion and each horizontal insertion, respectively, with the observer blind to the origin of the web. To avoid multiplication with zero, data averages were calculated after a value of one was added to a single increment of each of the nine measurements^42^.

To quantify the amount of contact pheromone components **1, 2** and **3** deposited on each web, the web was bundled up on a glass rod, transferred into a 1.5-mL glass vial and extracted overnight in 50 µL acetonitrile (99.9% HPLC-grade, Fisher Chemical, Ottawa, CA). A 2-µL aliquot was then analyzed on a Bruker maXis Impact Quadrupole Time-of-Flight HPLC/MS System. The system consisted of an Agilent 1200 HPLC fitted with a Spursil C_18_ column (30 mm × 3.0 mm, 3 µ; Dikma Technologies, Foothill Ranch, CA, USA) and a Bruker maXis Impact Ultra-High Resolution tandem TOF (UHR-Qq-TOF) mass spectrometer. The LC-MS was operated as follows: the mass spectrometer was set to positive electrospray ionisation (+ESI), with a gas temperature of 200 ºC and a gas flow of 9 L/min. The nebuliser was set to 4 bar and the capillary voltage to 4200 V. The column was eluted with a 0.4-mL/min flow of a solvent gradient, starting with 80% water and 20% acetonitrile, and ending with 100% of acetonitrile after 4 min. The peak shape of compounds in the extract was improved by adding 0.1% formic acid to the solvents. Contact pheromone components were quantified by comparing the area of their M + Na ion with that of the external standards **1** and **7**. Because a male spider courting on a web makes contact with single silk strands, the amount of contact pheromone components on each web was divided by its silk strand density (number of silk strands per web) to estimate the amount of contact pheromone components per silk strand in relation to the females’ age.

The airborne mate-attractant pheromone components (**4, 5, 6)** were quantified using **7** as a proxy for **4, 5**, and **6**, because **7** and the sum of **4, 5**, and **6** have equal stoichiometric quantities^37,42,49^. Because females can adjust the rate of pheromone hydrolysis and thus the dissemination of their mate-attractant pheromone components^42^, we used the ratio of **7** and **1**+**2**+**3**+**7** as an indicator of the females’ mate-call investment (essentially the attractiveness of their webs) in relation to their age.

The pH of web silk was measured because it determines the conversion rate of contact pheromone components to mate-attractant pheromone components^37,42,43,49^. After measuring the pH of 50 µL water (HPLC grade, EMD Millipore, Corp. Burlington, MA, USA) in a microvolume pH meter (LAQUAtwin pH 22 (Horiba, Kyoto, JP), a spider web was added, with the water being the conductor for the silk surface pH measurements^37,50^. Between measurements, the pH meter was rinsed with water and regularly re-calibrated, using pH 7 and pH 4 buffer solutions (Horiba, Kyoto, JP). The observer was blind to the origin of the silk.

### H3: Senescing females attract males as effectively as young females through deceptive signalling

Attraction of 50 naïve males to the web of a young female and an older female (50 pairs total) was tested in Y-tube olfactometers (main stem: 24 cm long, side arms 21 cm long, all 2.5 cm diameter) ^38,48^. The two females in each pair, with a mean age differential of 161 ± 4.7 days and with comparable body condition (Δ 0.015 ± 0.002), were placed on separate bamboo frames (see above) to build a web for three days. Each frame – together with the freshly-spun web and the female spider – was placed in a translucent oven bag (30 cm × 31 cm; Toppits, Mengen, DE), which was secured to the opening of a Y-tube side-arm. The olfactometer was lined with bamboo sticks to provide grip to the males. To initiate a bioassay, a single male was placed into a holding tube (26 cm long; 2.5 cm diameter) and allowed 2 min to acclimatize. The holding tube was then attached to the Y-tube olfactometer and air was drawn at 100 mL/min through the olfactometer. A response was recorded when the male entered one of the two oven bags within 5 min. The bamboo frames and oven bags were used only once, whereas all glassware was cleaned with hot soapy water and dried 3 h at 100 ºC in an oven.

### Statistical analyses

All data were analyzed statistically using R^51^. The relationship between the females’ lifetime reproductive output and several predictor variables was analyzed using a generalized linear model with tweedie distribution using the glmmTMB package. The predictor variables were female age, female and male condition, and the duration of male courtship and copulation, which were standardized using the scale argument. The relationship between web density (number of silk strands per web) and female age was analyzed using a GLM with tweedie distribution. The relationships between female age and the amount of female contact pheromone components, or pheromone hydrolysis product, were analyzed using individual GLMs. The amount of contact pheromone components, normalized on web density, was analyzed using a GLM with tweedie distribution. The web-normalized total amount of pheromone invested was analyzed using a negative binomial distribution (nbinom1), whereas the female condition model used the gaussian distribution. All models were tested to meet assumptions of residual uniformity and homogeneity of variance using the DHARMa package. A type III ANOVA of the car package was used to test for significance of predictor variables. Attraction data of males to webs of old and young females in olfactometer bioassays were analyzed using a binomial test, whereas male courtship duration and copulation duration were analyzed by GLMs with tweedie distribution, as described above.

## Results

### H1: Senescing females produce progressively fewer offspring

The lifetime reproductive output by females declined with progressive age (Fig. 2a), with the age of females at the time of mating being the only significant predictor for the number of spiderlings they produced (GLM, χ^2^ = 6.98, df = 1, p = 0.008). The indices for body condition of females and males did not contribute to the statistical model (both females and males: GLM, χ^2^ = 0.30, df = 1, p = 0.584, suppl. Figs. 2, 3). Similarly, neither the duration of male courtship nor the duration of copulation affected the number of spiderlings that females produced (GLM, courtship duration: χ^2^ = 0.13, df = 1, p = 0.723; copulation duration: χ^2^ = 0.10, df = 1, p = 0.758; suppl. Figs. 4, 5).

**Figure 2.**
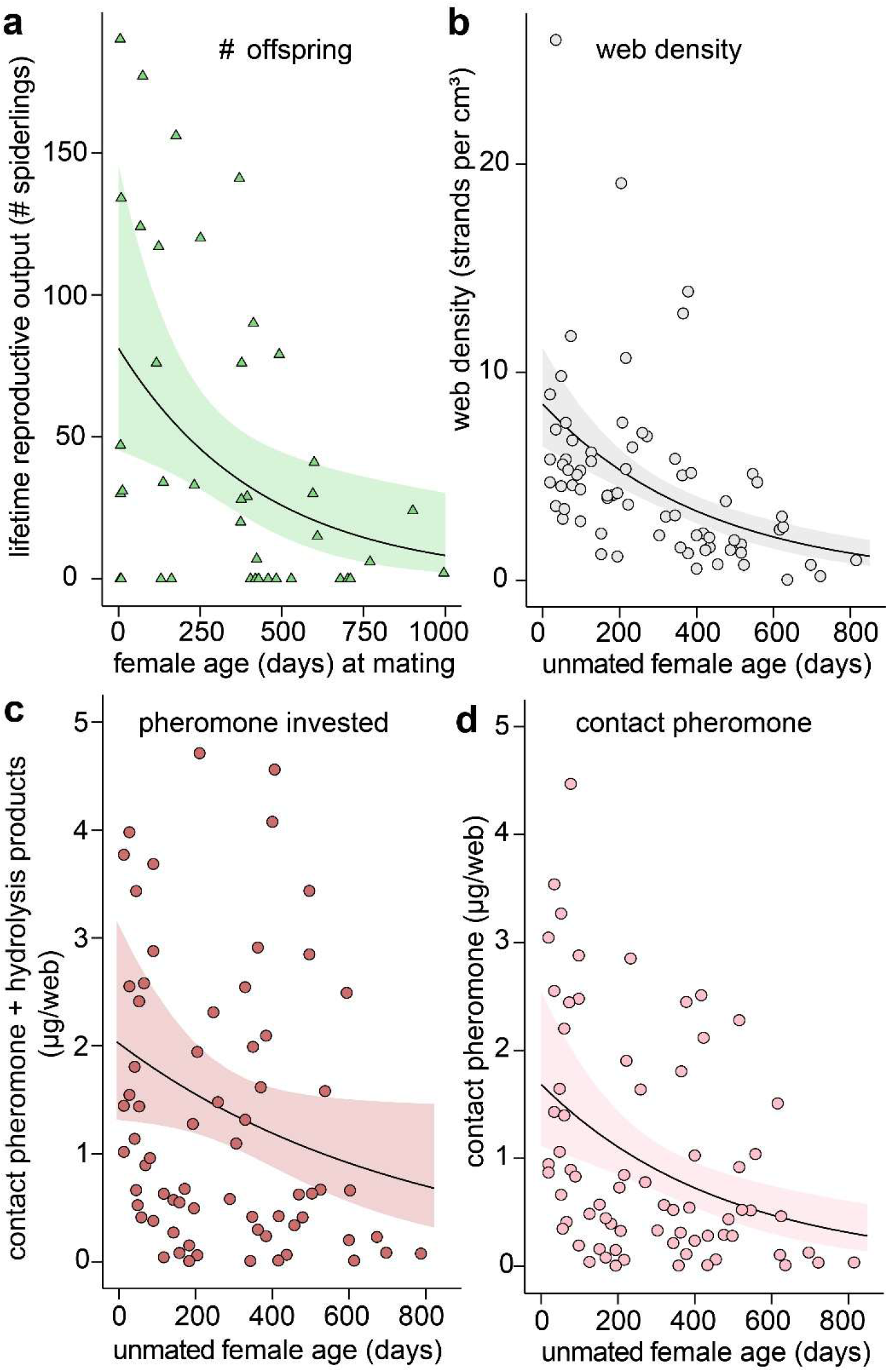
Effects of age, and age at mating, of female *Steadoda grossa* on web density, contact pheromone deposition, and lifetime reproductive output. (a) Lifetime reproductive output (# spiderlings produced) of females in relation to age at mating. (b) Mean number of silk strands per web (web density). (c) Mean amount of pheromone invested: contact pheromone components **1**+**2**+**3** (Figure 1) and their hydrolysis product **7** per web. (d) Mean amount of contact pheromone components (**1**+**2**+**3**) per web. (d) Solid lines show the mean and shaded areas 95% confidence intervals. All relationships shown in subpanels **a**-**d** are statistically significant (p<0.05, generalized linear models). Triangles represent data from female-male pairs, whereas circles represent web measurements.

### H2: Senescing females adjust their web density and sexual pheromone signals in accordance with their diminishing metabolic resources

Web density (the number of silk strands per web) declined with progressive female age (GLM, χ^2^ = 30.40, df = 1, p < 0.001, Fig. 2b). The amount of contact pheromone components **1**+**2**+**3** which females deposited on their web declined with female age (**1**+**2**+**3**, GLM: χ^2^ = 12.13, df = 1, p < 0.001, Fig. 2c), as did the amount of **1**+**2**+**3** together with their hydrolysis product **7** (**1**+**2**+**3**+**7**; GLM, χ^2^ = 3.942, df = 1, p = 0.047, Fig. 2c). In contrast, the amount of mate-attractant pheromone components – quantified using the amount of **7** on webs as a proxy – remained constant with female age (GLM, χ^2^ = 1.13, df = 1, p = 0.287, Fig. 3a). This phenomenon is explained by an age-dependent increase in mate-call investment (the dissemination of mate-attractant pheromone components from webs), as evident by an increasing conversion ratio of contact pheromone components to mate-attractant pheromone components with female age (GLM, χ^2^ = 11.33, df = 1, p < 0.001, Fig. 3b), despite the deposition of contact pheromone components declining with female age (Fig. 2c).

**Figure 3.**
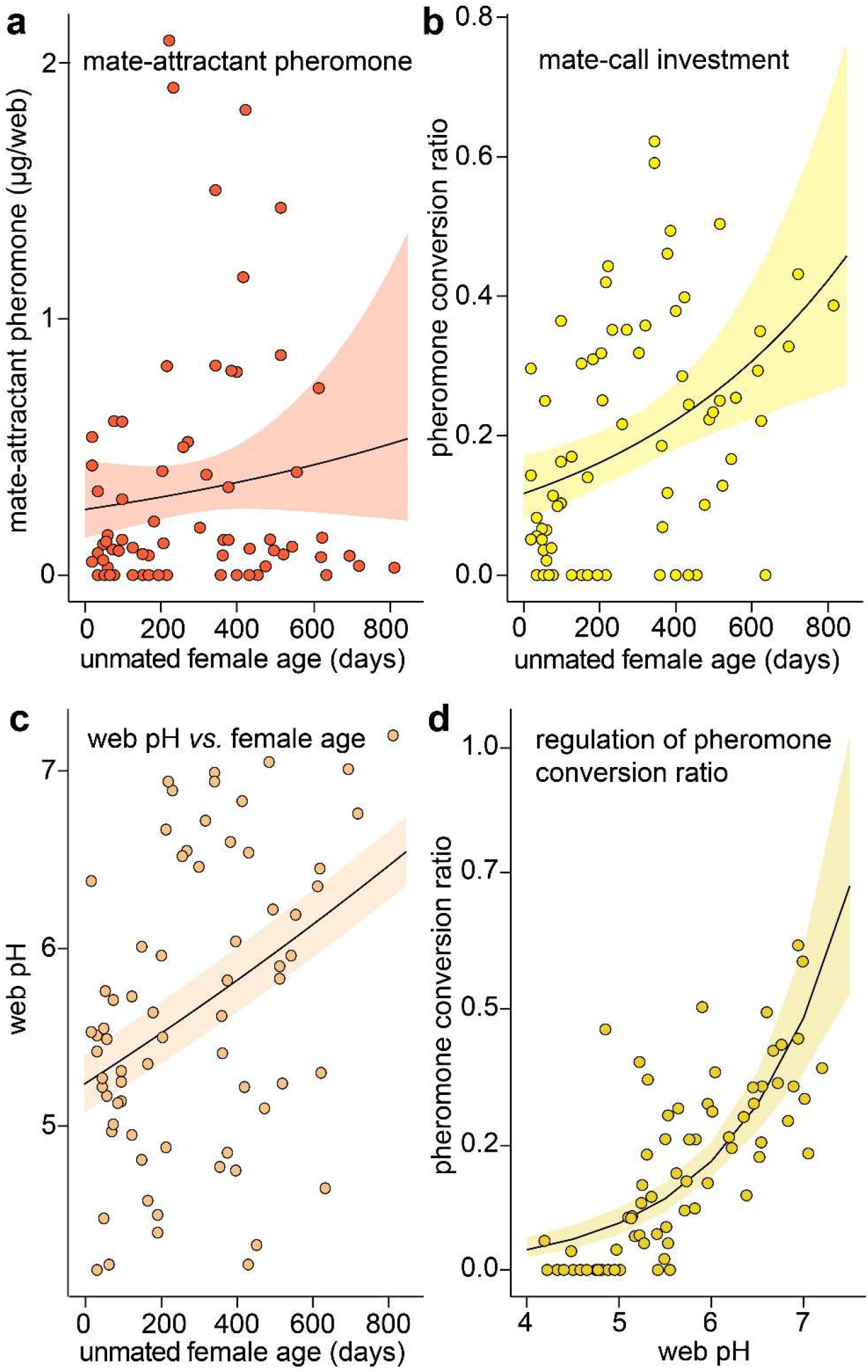
Age effects of unmated female *Steadoda grossa* on pheromone signals and web pH. (**a**) Amount of mate-attractant pheromone components (quantified via hydrolysis product **7** as a proxy; see Figure 1) released from webs of senescing females. (**b**) Conversion ratio of contact pheromone components to mate-attractant pheromone components (**7**/**1**+**2**+**3**+**7**) in relation to female age. (**c**) Relationship between web pH and age of unmated females. (**d**) Relationship between the contact pheromone conversion ratio (**7**/**1**+**2**+**3**+**7**) and web pH. Solid lines represent the mean and shaded areas 95% confidence intervals. Data are statistically significant (p< 0.05, generalized linear models) for subpanels **b, c** and **d** but not for subpanel **a**.

Silk surface pH increased with female age (Exp. 4, GLM, χ^2^ = 11.87, df = 1, p < 0.001, Fig. 3c) and was positively correlated with the hydrolysis rate of contact pheromone components to mate-attractant pheromone components (GLM, χ^2^ = 61.86, df = 1, p < 0.001, Fig. 3d)

### H3: Senescing females attract males as effectively as young females through deceptive signalling

In Y-tube olfactometers, 50 singly-tested males were equally often attracted to the web of a young female (good prospective mate) and an old female (poor prospective mate) in 50 pairs of females (binomial test, 21 *vs* 29 response ratio, p = 0.322, Fig. 4c). Progressive age of female (N = 41) shortened the time males engaged in courtship (GLM, χ^2^ = 4.71, df = 1, p = 0.029, Fig. 4a), consistent with decreasing amounts of contact pheromone components deposited by senescing females (Fig. 2d). Interestingly, female age did not affect the time females and males remained *in copula* (GLM, χ^2^ = 2.02, df = 1, p = 0.155, suppl. Fig. 8).

**Figure 4.**
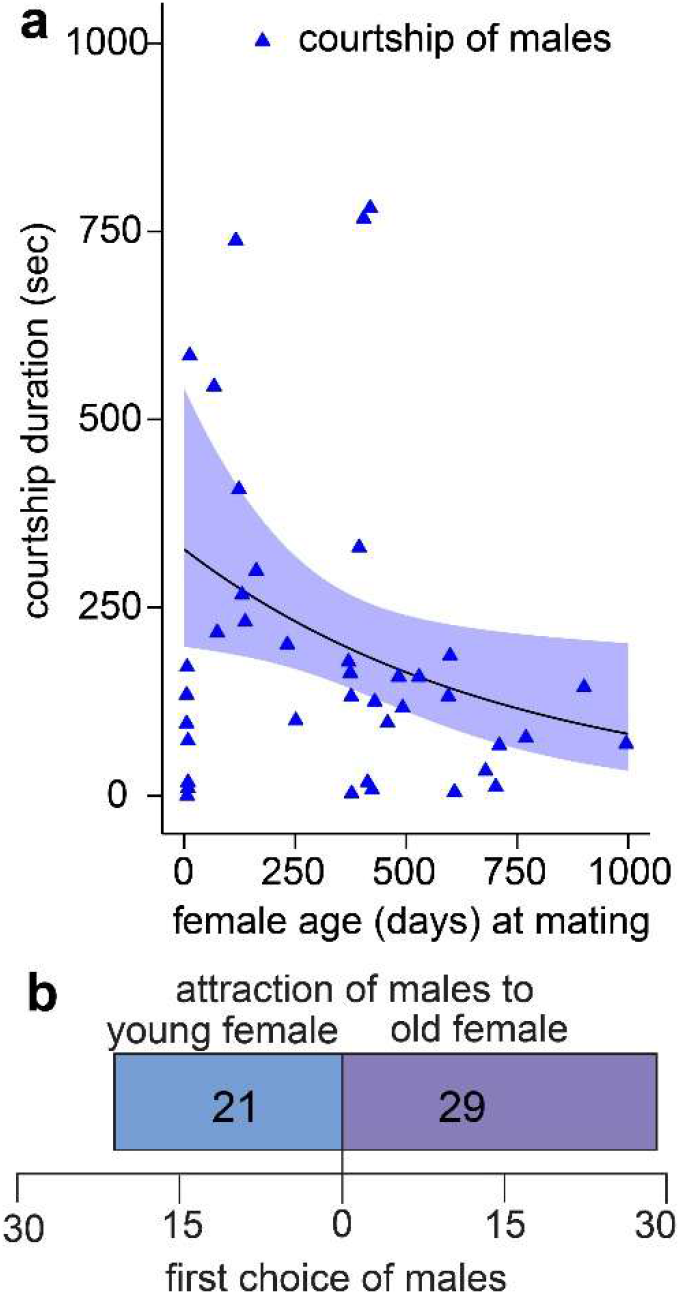
Behavioral responses of male *Steatoda grossa* to web-borne contact pheromone components and airborne mate-attractant pheromone components of conspecific unmated females. (a) Courtship duration of males on the web of females in relation to the age of females when they produced the web. The solid line represents the mean and the shaded area the 95% confidence interval (p = 0.029; generalized linear model). (b) Attraction of males in Y-tube olfactometers to mate-attractant pheromone components disseminating from the webs of young and old females.

## Discussion

Our data support the hypotheses that senescing female *S. grossa* produce progressively fewer offspring but adjust their sexual signals to remain attractive to mate-seeking males. Although old females are poor mates with limited residual reproductive value, they remained as attractive as young females by engaging in deceptive signalling at the expense of males who would sire more spiderlings by being attracted to, and mating with, young females.

The decline in residual reproductive output by female *S. grossa* in relation to their age at mating (Fig. 2a) is common among animals^20,52^, including spiders. For example, aging unmated brown widow spiders, *Latrodectus geometricus*, produce progressively fewer viable eggs^53^. Stronger attraction of male *L. geometricus* to older unmated females than to younger females, and greater propensity of males to mate with older females, are thought to be caused by deceptive chemical signalling by older females (Waner et al. 2018). Our study reveals the mechanisms underlying deceptive signalling in *S. grossa* (see below), a close phylogenetic relative of *L. geometricus* with a similar pheromone system (reviewed in^54^). The residual reproductive output of *S. grossa* females was linked to their age, with other factors such as body condition of mating partners not playing a role. Because copulation duration did not predict the number of spiderlings produced by females, it follows that sperm is likely transferred early during copulation^55^. However, the effects of copulation duration and repeated copulations on offspring production should be investigated further^56^.

*Steatoda grossa* males mating with old females of low residual reproductive value (poor mates) not only incur reproductive fitness costs but also metabolic and opportunity costs, and they risk being preyed upon during mate search^23,24,57^. Upon arrival on a web, male *S. grossa* engage in costly courtship by depositing metabolically-expensive proteinaceous silk on the female’s web^48^. All data combined support the conclusion that old unmated females are poor mates that resort to deceptive signalling to meet their need of securing a mate.

The terminal investment strategy predicts that aging unmated signalers with an ever-increasing need for mates amplify their efforts to attract a mate, potentially misrepresenting their residual reproductive value through deceptive signalling (Fig. 1a)^21,22^. In our study, old unmated *S. grossa* females remained as attractive to males as young unmated females (Fig. 4) despite their lower residual reproductive value (Fig. 2a), aligning with the ‘need-based’ deceptive signaling concept (Fig. 1a). Even though old females deposited less contact pheromone components on their web (Fig. 2c), they remained attractive to males by accelerating the conversion of contact pheromone components to mate-attractant pheromone components (Figs. 1b+3b). The accelerated dissemination of mate-attractant pheromone components from webs of senescing females may be attributed to a progressively increasing web pH (Fig. 3c) which, in turn, likely affected the enzymatic activity of a carboxyl ester hydrolase (CEH) present on *S. grossa* webs^37,58^.

Assuming that it is this CEH that hydrolyzes the contact pheromone components at their ester bond, the spiders would be able to regulate the enzym’s activity by adjusting their silk’s pH^37,41^. If this assumption is correct, senescing unmated *S. grossa* females seem to retain the attractiveness of their webs by increasing their web’s pH, and thus the dissemination rate of mate-attractant pheromone components.

Whether old(er) female spiders remain attractive to conspecific males is dependent upon their signalling strategy^36^. Monandrous female spiders seem to seize signalling and loose attractiveness to males upon mating^19,36,59^, whereas senescing polyandrous female spiders remain attractive, or become more attractive, regardless of their mating status^60^. Females of *S. grossa* are considered polyandrous because they re-mate, although not as likely as unmated females^61^. Sustained or even increased attractiveness of senescing polyandrous female spiders is well documented (reviewed in^36^), but the disconnect between the attractiveness of old females and their residual reproductive value has rarely been noted^53,62^. Moreover, reproductive senescence in arthropods has rarely been studied^8,20,52^ as have the underlying sexual signals that senescing unmated females produce^8,52,59^. Unmated female wasp spiders, *Argiope bruennichi*, increase signaling throughout their 14-day mating season and then deposit eggs that are fertilized or not^19,59^. Most female *A. bruennichi* get mated already while molting to adults^63^. Generally, remaining unmated for long is thought to be costly^22^ but empirical field records of old unmated females are rare. According to a field survey, only few female *L. hesperus* (4%) remained unmated at the end of the mating season^64^ but these female could still mate outside the peak mating season^54^. Long-lived females with multiple mating seasons incur less severe costs for remaining unmated, but their signalling strategy likely evolved mitigating the risk of remaining unmated^22^.

Chemical communication signals of arthropods other than insects remain largely unexplored^3,32–34^, and sexual signals of males are more often studied than those of females^28,30,31^. Senescent adult moths spent more time pheromone signalling, although the actual release of pheromone seems to underly species-specific constraints^8^. Similarly, the pheromone-based attractiveness of female mealybugs, *Planococcus kraunhiae*, increased with ageing, although the females’ reproductive performance was in decline, suggesting a sexual conflict within a pheromone signalling system, where females benefit through deceptive signals of fertility at the expense of males^65^. Generally, the onset of terminal investment to attract a mate may have a dynamic threshold because not only the signaller’s age but multiple other factors such as immune challenge may affect the trade-off benefit of this risky signalling strategy^66^. It is noteworthy that sexual signals remain evolutionary stable only if cheaters – the cohort of senescent spiders in our study – are rare, and thus males are rarely selected to avoid terminal-investment signals^22^. It follows that sexual signals, on average, are honest^67^.

Senescing female *S. grossa* produced fewer silk strands per web (Fig. 2b) and deposited less contact pheromone on their webs (Fig. 2c+d), honestly reflecting their declining residual reproductive value, as predicted by the ‘state-based’ signaling concept (Fig. 1a). Males courting on the web of females apparently integrated this honest information about the females’ physiological state because they adjusted the duration of their courtship in accordance with female age (Fig. 4). Apparently, males assessed the overall amount of contact pheromone on the web rather than pheromone amounts present on individual silk strands which did not decline with female age (Fig. S7). That old(er) females produced lower-density webs is likely due to the intrinsic costs of silk. Web-spinning is nutrient-demanding, requiring approximately 4.5 calories for each milligram of silk fiber^68^. The time spent building a web also incurs opportunity costs, detracting from prey capture activities^69^. The metabolic costs of silk are further reflected in observations that web-building spiders save silk-production costs by taking over existing webs^70,71^. The nitrogen-containing contact pheromone components of widow spiders are thought to incur additional nutritional costs, serving as a mechanism that ensures signal honesty^72^. Although old females deceptively lured males to their webs (see above), they presented honest signals at their webs, likely owing to inherent metabolic demands of silk and contact pheromone production^67^. These honest signals, however, apparently did not diminish the females’ chances of mating, even though males mating with an old female, rather than a young female, would sire fewer spiderlings (Fig. 2a). The males’ reproductive fitness benefit of mating with an old(er) female may outweigh the costs of rejecting an old female and resuming mate search for a young female, a search which may or may not be successful.

Pheromone signals of spiders incur metabolic production costs and enhance predation risk for the signaler because predators eavesdrop on these signals^19^. In our study with *S. grossa*, pheromone production costs and ecological costs can be disentangled. Females bear metabolic expenses only for the production of courtship-inducing contact pheromone components^37^, and only the airborne mate-attractant pheromone components are susceptible to eavesdropping by specialized predators^73,74^. That contact pheromone deposition on webs declined with female age (Fig. 2c) is indicative of high pheromone production costs that old(er) females seem unable to afford. As Baruffaldi and Andrade (2015) argued, the nitrogen-rich pheromone components, derived from amino acid metabolism, incur nutritional costs that only high-quality females may be able to afford^72^. This concept of costly pheromone production deviates from presumptions that pheromone biosynthesis is inexpensive^5,9–12^. Moreover, studies on moths and ants attribute substantial costs to pheromone production, including accelerated mortality of the pheromone signaller^13–16^. However, a general statement about pheromone production cost might be difficult because biosynthetic pathways are diverse^75^ and energy costs may differ with pathway and evolutionary origin of signals. Signals that evolved from metabolic waste may incur low production costs, whereas pheromones originating from energetically expensive precursors likely incur higher costs. To test this prediction, we envision comparative studies that examine the metabolic costs of various pheromone biosynthetic pathways.

Ecological costs of pheromone signals arise when predators or parasitoids intercept these signals for locating prey or hosts. Airborne pheromones are considered at higher risk of being intercepted than colorful visual signals, according to a meta-analysis across taxa^11^. Specialized predators of spiders, such as mud dauber wasps, locate their prey by intercepting prey scent^76^. Some of these wasps may hunt any spider they encounter but prey inventories of these wasps often revealed species-specific and female-biased preferences for prey^73^, supporting the concept of prey pheromone interception^19,42^. Because our study on *S. grossa* took place in laboratory settings and predators were not available, we could not test attraction of predators to signalling females. Potential trade-offs between pheromone signalling and predation risk should be investigated in future studies, especially in light of terminal investments by the signaller. In general, more chemo-ecological studies of non-model taxa are needed to increase our understanding of complex communication systems^34^.

In conclusion, our study revealed the complex and evolving communication system of *S. grossa* females that is shaped by need-dependent and state-dependent constraints of senescing females (Fig. 1a). High pheromone production costs seem to enforce state-dependent honest signalling, whereas the ever-increasing need for mates experienced by senescing females invokes deceptive signalling, resulting in the attraction of males to old females that are poor mates with limited residual reproductive value. Even though males mating with an old female, rather than a young female, sire fewer spiderlings, the males’ reproductive fitness benefit of mating with an old(er) female may still outweigh the costs of rejecting an old female and resuming mate search for a young female, a search which may, or may not, be successful.

## Acknowledgements

We thank Gabriele Uhl for valuable comments on previous versions of this manuscript. A.F. was supported by Graduate Fellowships from SFU, the H.R. McCarthy Bursary, and an Alexander Graham Bell Scholarship from the Natural Sciences and Engineering Research Council of Canada (NSERC). The project was funded by an NSERC-Industrial Research Chair to G.G., with BASF Canada Inc. and Scotts Canada Ltd. as the industrial sponsors. The research was further supported by a Student Research Grant from the American Arachnological Society to A.F.. The funders had no role in study design, data collection and analysis, decision to publish, or preparation of the manuscript. Open Access funding enabled and organized by Projekt DEAL.

## Author contributions

Conceptualization, A.F.; Methodology, A.F.; Investigation, A.F., A.C.R.T., and B.C.; Data Curation, A.F.; Writing – Original Draft, A.F; Writing – Review & Editing, G.G. Funding Acquisition, G.G., A.F.; Resources, G.G., A.F.; Supervision, A.F.

## Data availability statement

Source data are available in Supplementary Data. All other data supporting the findings of this study are available within the paper and its Supplementary Information.

## Data availability statement

Code to analyse the data is available as Supplementary Data.

