## Supplementary material for "Senescent widow spiders are poor mates but remain attractive to mate-seeking males through deceptive signalling": Suppl. Figures

Supplementary figures

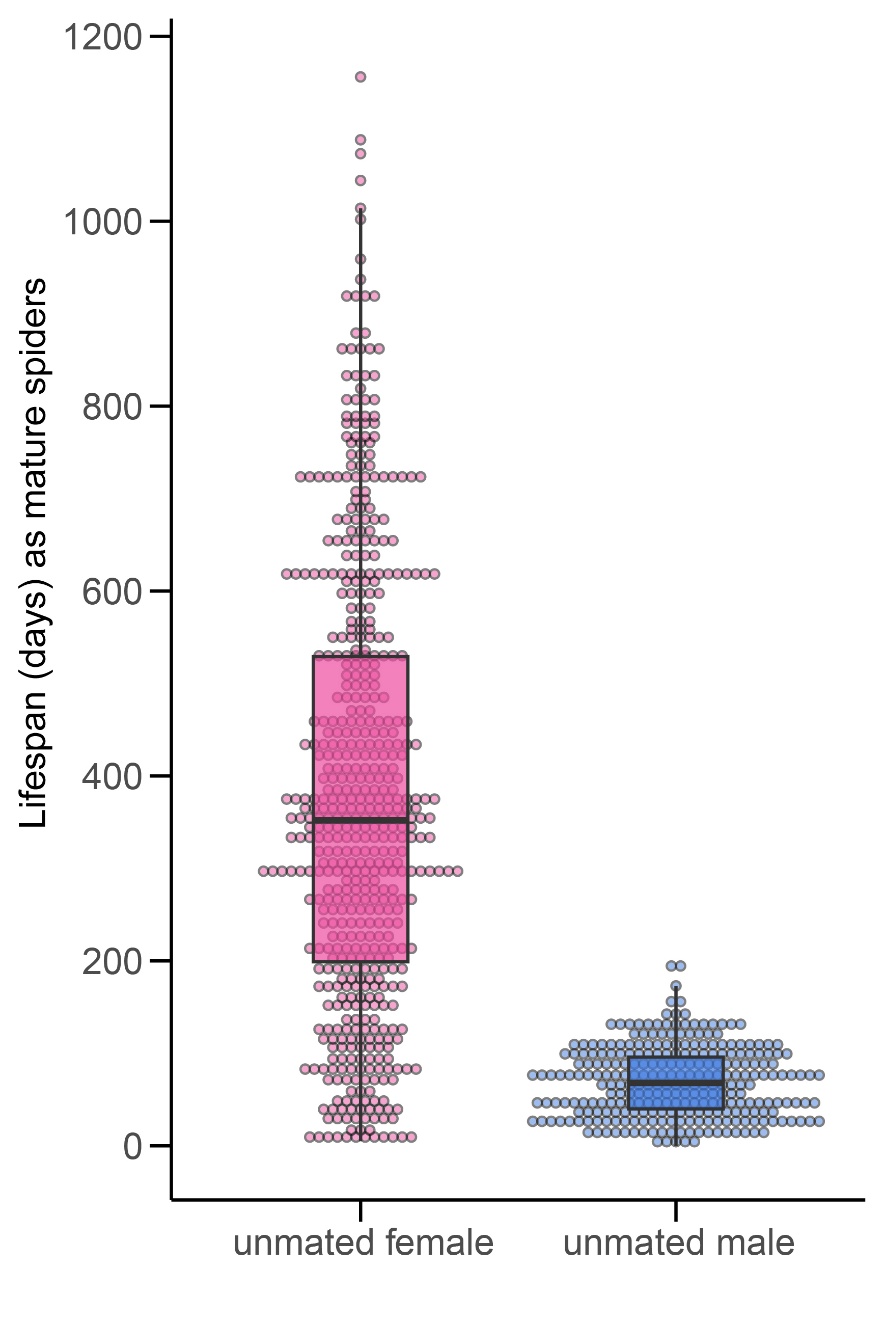

Figure S1: Lifespan of sexually mature unmated Steatoda grossa females and males (N_Females_ = 553, N_Males_ = 277). Solid circles indicate the lifespan of individual spiders, the central line in each box represents the median, and the box top and bottom represent the third and first quartiles, respectively.

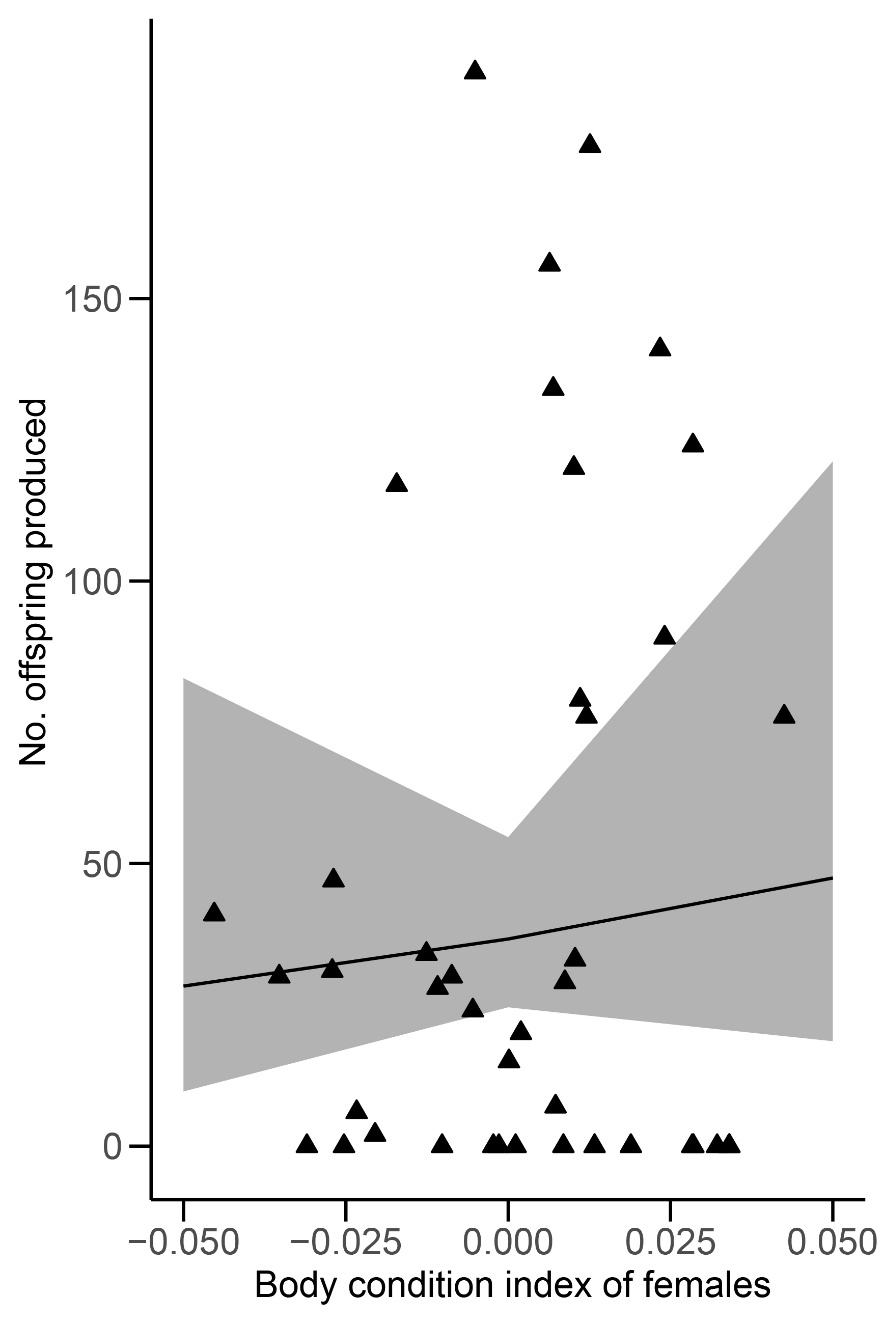

Figure S2: Relationship between the body condition index of female Steatoda grossa in 41 female-male pairs and the number (No.) of viable offspring they produced. Body condition indices were calculated as the residuals of the relationship between body mass and body size (tibia-patella length) (Fischer et al. 2020). Triangles represent the data of individual spiders, whereas the line and shaded area represent the mean and 95% confidence interval, respectively (generalized linear model, p>0.05).

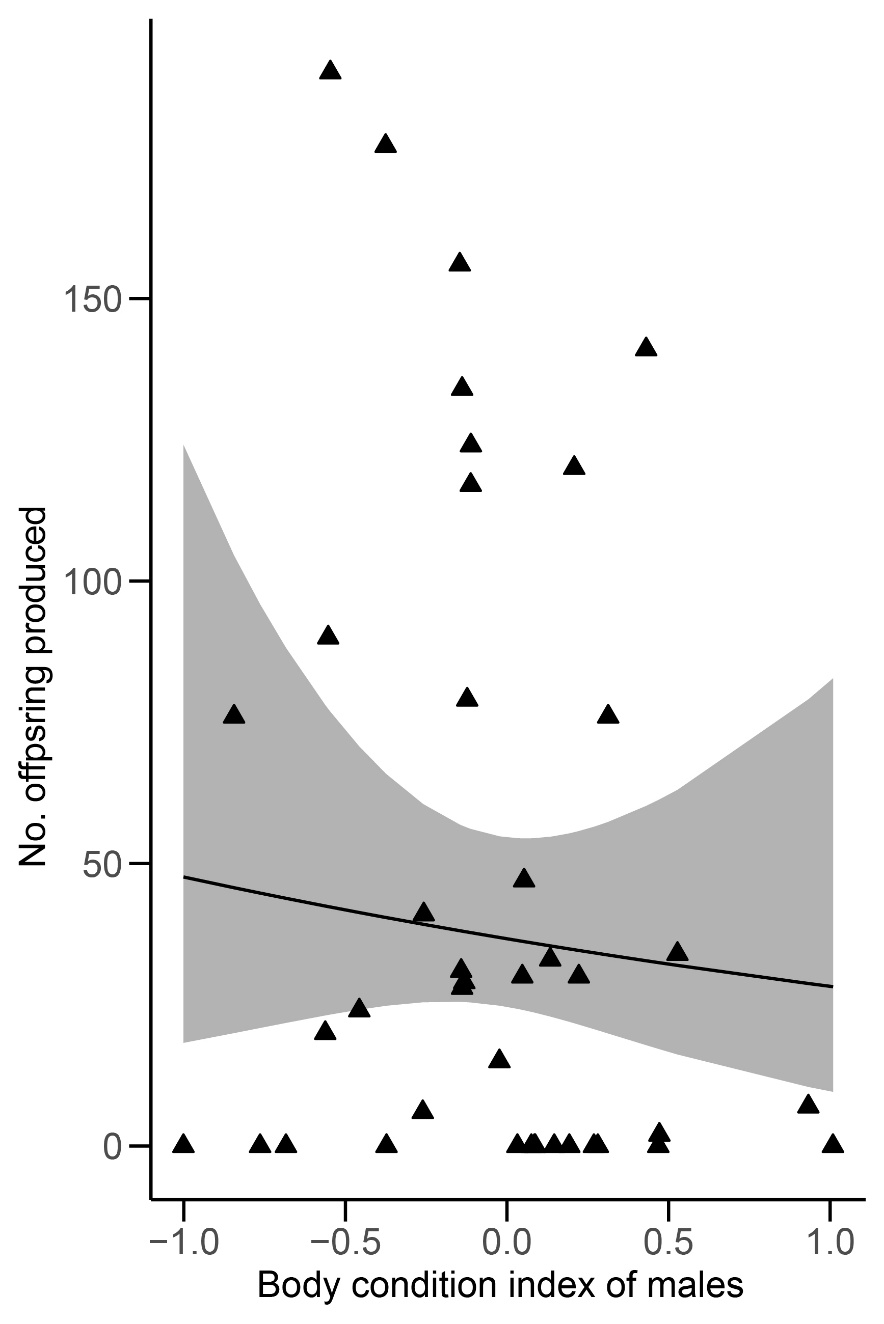

Figure S3: Relationship between the body condition index of male Steatoda grossa in 41 male-female pairs and the number (No.) of viable offspring females produced. Body condition indices were calculated as the relationship between residuals of body mass and body size (tibia-patella length) (Fischer et al. 2020). Triangles represent the data of individual spiders, whereas the line and shaded area represent the mean and 95% confidence interval, respectively (generalized linear model, p>0.05).

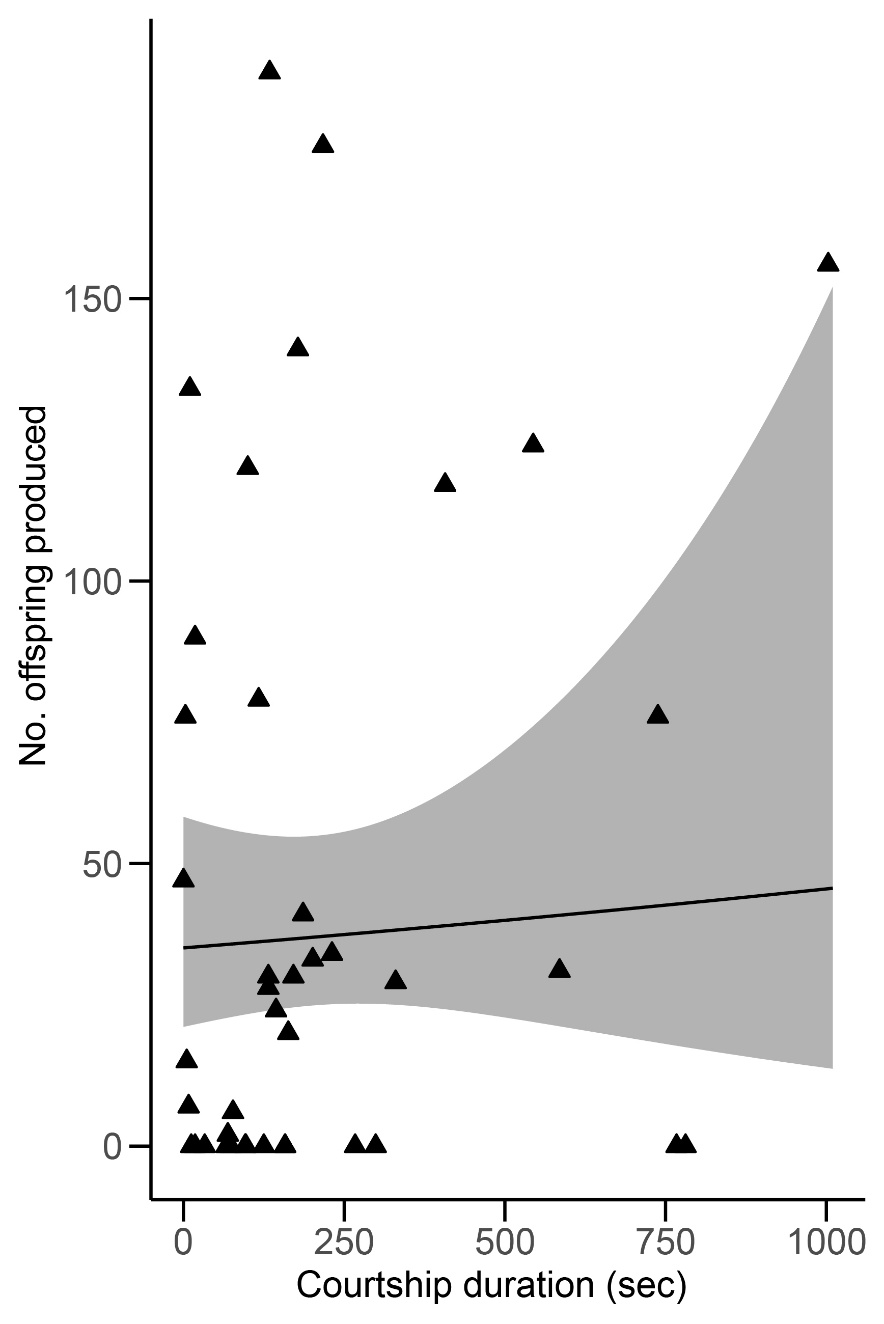

Figure S4: Relationship between the courtship duration of male Steatoda grossa in 41 female-male pairs and the number (No.) of viable offspring females produced. Triangles represent the data of individual spiders, whereas the line and shaded area represent the mean and 95% confidence interval, respectively (generalized linear model, p>0.05).

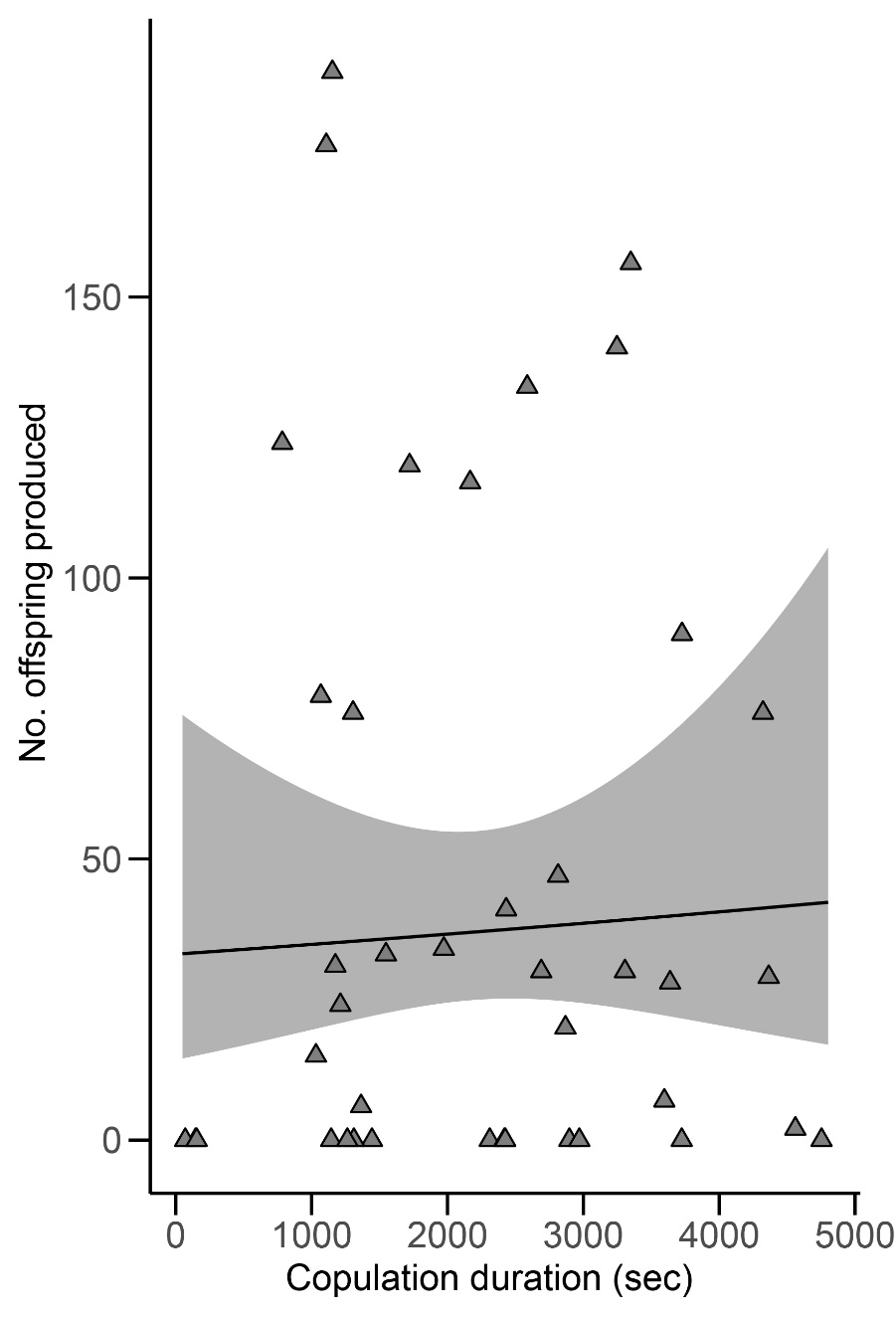

Figure S5: Relationship between copulation duration in 41 female-male Steatoda grossa pairs and the number (No.) of viable offspring females produced. Triangles represent the data of individual spiders, whereas the line and shaded area represent the mean and 95% confidence interval, respectively (generalized linear model, p>0.05).

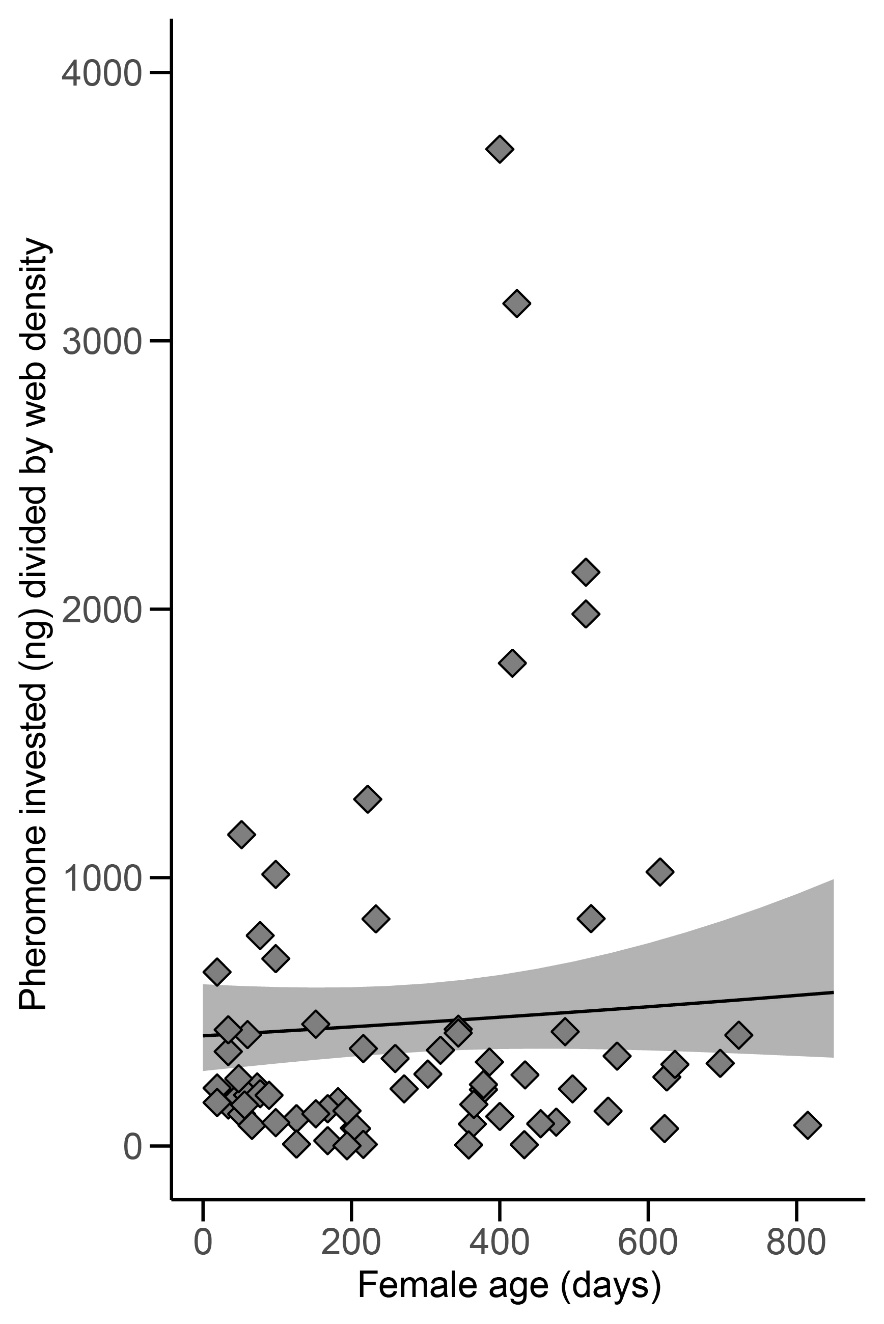

Figure S6: Relationship between the age of Steatoda grossa females (N = 41) and the amount of pheromone (contact pheromone components plus hydrolysis product) they invested (deposited) on their web divided by web density (silk strands per web). Rhombi represent the data of individual spiders, whereas the line and shaded area represent the mean and 95% confidence interval, respectively (generalized linear model, p>0.05).

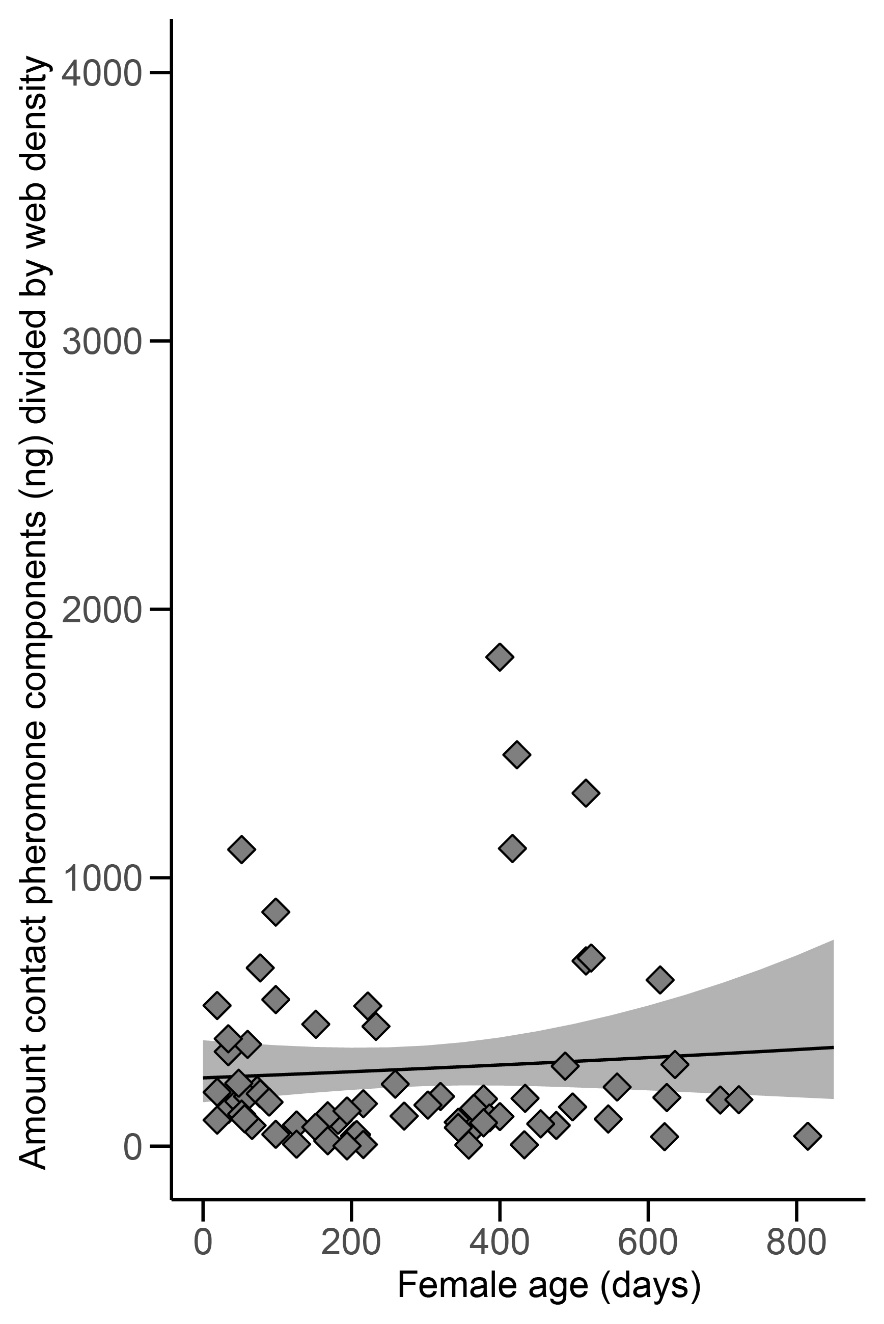

Figure S7: Relationship between the age of Steatoda grossa females (N = 41) and the amount of contact pheromone components they invested (deposited) on their web divided by web density (silk strands per web). Rhombi represent the data of individual spiders, whereas the line and shaded area represent the mean and 95% confidence interval, respectively (generalized linear model, p>0.05).

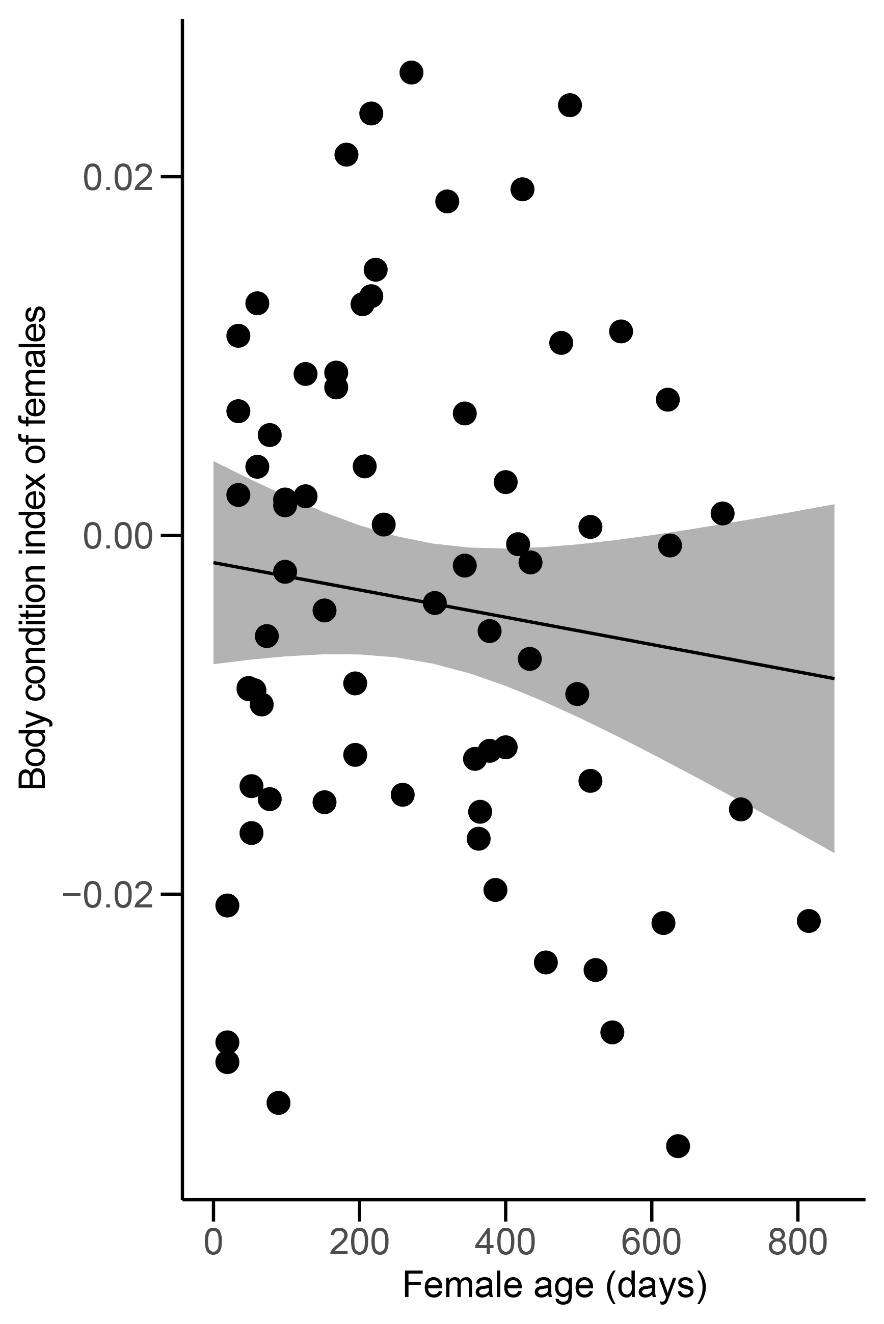

Figure S8: Relationship between the age of Steatoda grossa females (N = 41) and their body condition. Body condition indices were calculated as the residuals of the relationship between body mass and body size (tibia-patella length) (Fischer et al. 2020). Solid circles represent the data of individual spiders, whereas the line and shaded area represent the mean and 95% confidence interval, respectively (generalized linear model, p>0.05).

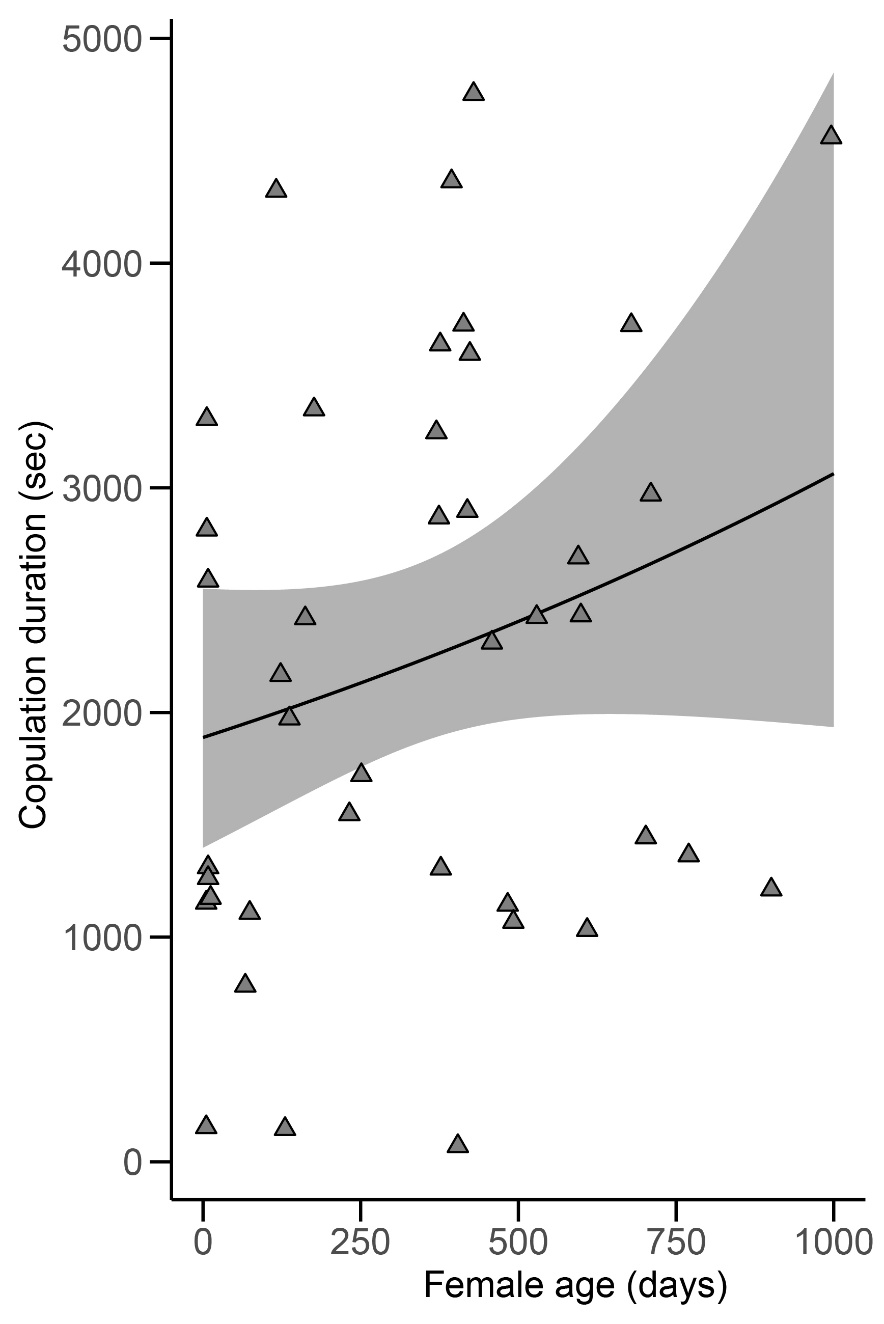

Figure S9: Relationship between the age of Steatoda grossa females in 41 female-male pairs and the duration of copulation. Triangles represent the data of individual spiders, whereas the line and shaded area represent the mean and 95% confidence interval, respectively (generalized linear model, p>0.05).
